# CD4^+^ T cells from chronic Chagas disease patients with different degrees of cardiac compromise exhibit distinct expression patterns of inhibitory receptors TIGIT, Tim-3 and Lag-3

**DOI:** 10.1101/694729

**Authors:** Paula B. Alcaraz, Magali C. Girard, M. Paula Beati, Raul Chadi, Marisa Fernandez, Yolanda Hernandez, Karina A. Gómez, Gonzalo R. Acevedo

## Abstract

T cells are central to adaptive immune response against *T. cruzi* infection. In the chronic stage of Chagas disease, circulating parasite-specific memory T cells show reduced functionality and increased expression of inhibitory receptors, possibly as a result of persistent antigenic stimulation. This exhausted phenotype has been linked to progression of cardiac pathology while, contrariwise, the presence of polyfunctional T cells shows association with therapeutic success and more efficient control of infection. Given this, we hypothesized that inhibitory receptors TIGIT, Tim-3 and Lag-3 may be involved in immune modulation of anti-*T. cruzi* T cell response, and therefore may play a role in the containment or the unleashing of inflammatory phenomena that ultimately lead to tissue damage and pathology. In this preliminary study, we assess the frequency of CD4^+^ T cells expressing each of these receptors and their relation to cellular activation. Samples from chronic Chagas disease patients with different degrees of cardiac compromise, and non-infected donors were analyzed under different stimulation conditions. Our results show that the frequency of TIGIT^+^ CD4^+^ T cells is increased in Chagas patients, while Tim-3^+^ cells are more abundant in patients with signs of cardiac alterations. In addition, the frequency of Lag-3^+^ cells increases in non-activated CD4^+^ T cells from Chagas patients without demonstrable cardiopathy upon pathogen-specific *in vitro* antigenic stimulation.

## Introduction

Chronic infections are one of the circumstances under which T cells encounter the challenge of appropriately regulating the immune response in persistent presence of antigenic stimulus. In such settings, a process known as T cell exhaustion takes place, by which these cells acquire features believed to hamper sterile immunity, such as hierarchical loss of effector functions, altered gene expression regulation and metabolic disarrangements^1–3^. The increased and/or sustained expression of inhibitory receptors on the cellular surface of T cells is another of these characteristics, and is often regarded as the hallmark of cell exhaustion^3,4^.

In chronic infection with *Trypanosoma cruzi*, i.e. chronic Chagas disease, evidence of exhausted T cells in circulation has accumulated during the last decades^5^. In fact, the frequency of cells with reduced effector capabilities has been directly correlated to cardiac compromise^6,7^. The association of therapeutic success of benznidazole with a reduction of T cells with an exhausted profile in the circulation of treated subjects^8,9^ and the observation of a richer response profile in serodiscordant subjects^10^ further support the relevance of persistent antigen-specific activation for ineffective adaptive immunity against the parasite.

Although PD-1 and CTLA-4 were the pioneer inhibitory receptors to be studied in relation to T cell exhaustion, other molecules were demonstrated to complement their function in a lower hierarchical level, apparently enabling the fine-tuning of T cell inhibition^2,4,11^. Among these molecules, TIGIT, Tim-3 and Lag-3 have been found to be implicated in chronic viral infections, and arise as promising targets for immune-enhancing therapies^4,11^. Their role in chronic Chagas disease is yet to be investigated.

The study of inhibitory receptors and their relationship with T cell exhaustion requires the analysis of T cell activation upon antigen specific stimulation, but the heterogeneity of response profiles displayed by the CD4^+^ T cell subset is a hurdle to the measurement of overall activation. This has been addressed by several research groups, resulting in the development of flow cytometry techniques based on the detection of molecules, globally referred to as activation induced markers (AIM)^12–14^. Here, we take advantage of this methodology to characterize the expression of TIGIT, Tim-3 and Lag-3 in peripheral blood CD4^+^ T cells from chronic Chagas disease patients and their relation to *T. cruzi*-specific and non-antigen-specific activation.

## Donors, materials and methods

### Subjects inclusion and blood sample collection

Blood samples were collected from non-infected donors and patients with chronic Chagas disease after the nature of this study was explained to them and written consent was given, in accordance with the guidelines of the protocol approved by the Medical Ethics Committee of the and the Hospital General de Agudos “Dr.Ignacio Pirovano” and the Instituto Nacional de Parasitologia “Dr.M.Fatala Chaben”. The sample collection protocol followed the tenets of the declaration of Helsinki. All subjects were of age at the time of sample collection. The study population included 20 patients with chronic Chagas disease with at least 2 reactive serological tests, who were categorized, according to Kuschnir’s classification^15^ into groups A (without demonstrable cardiac pathology, Kuschnir class 0, n=10) or C (with signs of cardiac alterations, Kuschnir class 1 or 2, n=10). Ten donors were included as well, with negative serology for *T. cruzi* infection.

Samples consisted in 35 to 50 ml peripheral venous blood, collected in EDTA anti-coagulated tubes and processed up to 4 h after collection. Peripheral blood mononuclear cells (PBMC) were isolated by centrifugation (400 ×g, room temperature, 40 min) in a Ficoll-Paque gradient medium (GE Healthcare Bio-Sciences, Uppsala, Sweden), quantified by manual count in Neubauer chamber, and aliquoted in fetal bovine serum (FBS; Natocor, Córdoba, Argentina) with 10% v/v DMSO to be cryopreserved in liquid nitrogen until used.

### Parasite lysate

*T. cruzi* trypomastigote/amastigote lysate was prepared from VERO cells infected with Sylvio strain (Discrete Typing Unit TcI^16^) parasites (MOI 3:1) supernatants, as described elsewhere^17^. After lysis, the suspension was filter-sterilized through a 0.2 µm pore-size membrane, aliquoted, and stored at −80 °C until use.

### Cell culture and stimulation

For antigen stimulation, 3×10^6^ PBMC from each subject were seeded in 6 wells from a 96-well U-bottom plates and cultured in assay medium alone or with 10 µg/ml *T. cruzi* lysate, or 5 µg/ml phytohaemagglutinin (PHA, Sigma, St Louis, MO, USA). Cells were incubated at 37°C in a humidified, 5% CO_2_ atmosphere for 18h.

### Flow cytometry analysis of T cells

After culture, cells were centrifuged at 400 ×g for 10 min at room temperature (RT) and supernatants were discarded. Next, they were washed with PBS by centrifugation at 700 ×g during 3 min at RT, transferred into a 96-well V-bottom plate and resuspended in 25 µl of staining solution, containing the antibodies detailed in Table 1, diluted in 1X live/dead fixable viability dye (Zombie-Aqua, Biolegend, San Diego, CA, USA). After staining for 30 min at RT, cells were washed with PBS, fixed with Fixation Buffer (Biolegend) for 20 min at RT in the dark and washed with PBS. Isotype control stained samples were used to set the cut point values for each marker. A minimum of 5×10^5^ events within the lymphocyte population were acquired in a FACSCanto II (BD Biosciences, San Diego, CA, USA) flow cytometer using FACS Diva Software (BD Biosciences). Flow cytometry analysis was carried out with the program FlowJo (FlowJo LLC, Ashland, OR, USA). All antibodies were used at optimal concentrations determined by previous titration experiments.

**Table 1.**
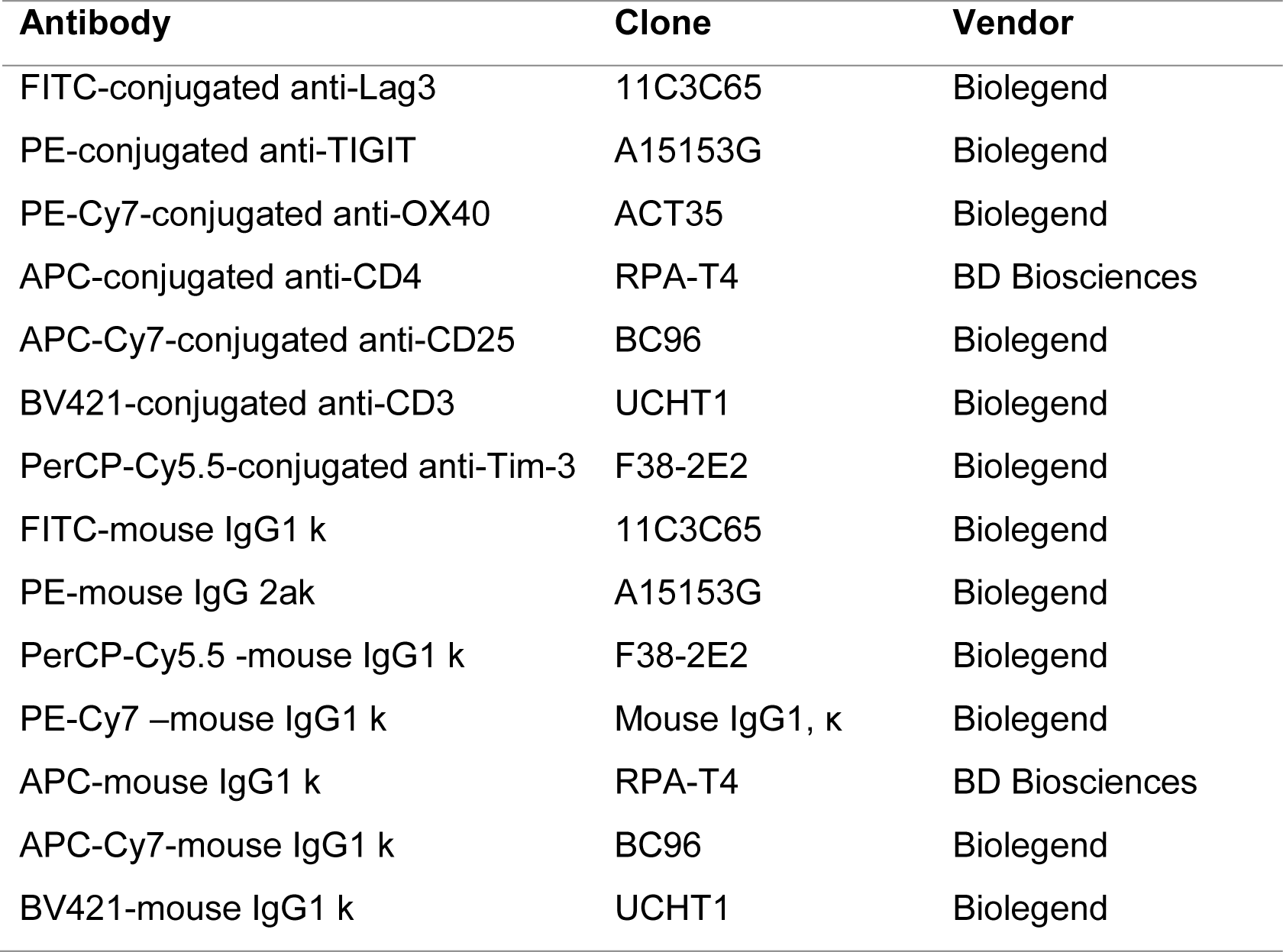
Fluorescent-labeled antibodies and isotype controls used for flow cytometry experiments.

| Antibody | Clone | Vendor |
| --- | --- | --- |
| FITC-conjugated anti-Lag3 | 11C3C65 | Biolegend |
| PE-conjugated anti-TIGIT | A15153G | Biolegend |
| PE-Cy7-conjugated anti-OX40 | ACT35 | Biolegend |
| APC-conjugated anti-CD4 | RPA-T4 | BD Biosciences |
| APC-Cy7-conjugated anti-CD25 | BC96 | Biolegend |
| BV421-conjugated anti-CD3 | UCHT1 | Biolegend |
| PerCP-Cy5.5-conjugated anti-Tim-3 | F38-2E2 | Biolegend |
| FITC-mouse IgG1 k | 11C3C65 | Biolegend |
| PE-mouse IgG 2ak | A15153G | Biolegend |
| PerCP-Cy5.5 -mouse IgG1 k | F38-2E2 | Biolegend |
| PE-Cy7 -mouse IgG1 k | Mouse IgG1, $\kappa$ | Biolegend |
| APC-mouse IgG1 k | RPA-T4 | BD Biosciences |
| APC-Cy7-mouse IgG1 k | BC96 | Biolegend |
| BV421-mouse IgG1 k | UCHT1 | Biolegend |

### Statistical analysis

The effect of the stimulation condition within each group was analyzed by Friedman’s test followed by multiple comparisons by pairwise Wilcoxon’s test, with data points paired by patient. The effect of groups within each stimulation condition was evaluated by Kruskal-Wallis’s test, followed by multiple comparisons by Dunn’s test. Significance was considered using α=0.05. For both multiple comparisons methods, *p* values were adjusted using the Benjamini-Yekutieli method. All statistical analysis methods were implemented using open source R packages stats^18^ and dunn.test^19^.

## Results

### Ox40 and CD25 are useful as activation markers in T. cruzi specific CD4+ T cell response

Seeking to determine whether the AIM assay proposed by Reiss et al.^13^ is useful for the measurement of CD4^+^ T cell activation against *T. cruzi* antigens, PBMC from non-infected individuals (group NI), and chronic Chagas disease patients with (group C) and without (group A) cardiac compromise were stimulated *in vitro* with parasite lysate to evaluate pathogen-specific response, and with PHA to assess overall, non-specific activation. Cells were stained and analyzed by flow cytometry. The gating strategy used for this analysis is represented in **Figure 1A** and **B**.

**Figure 1.**
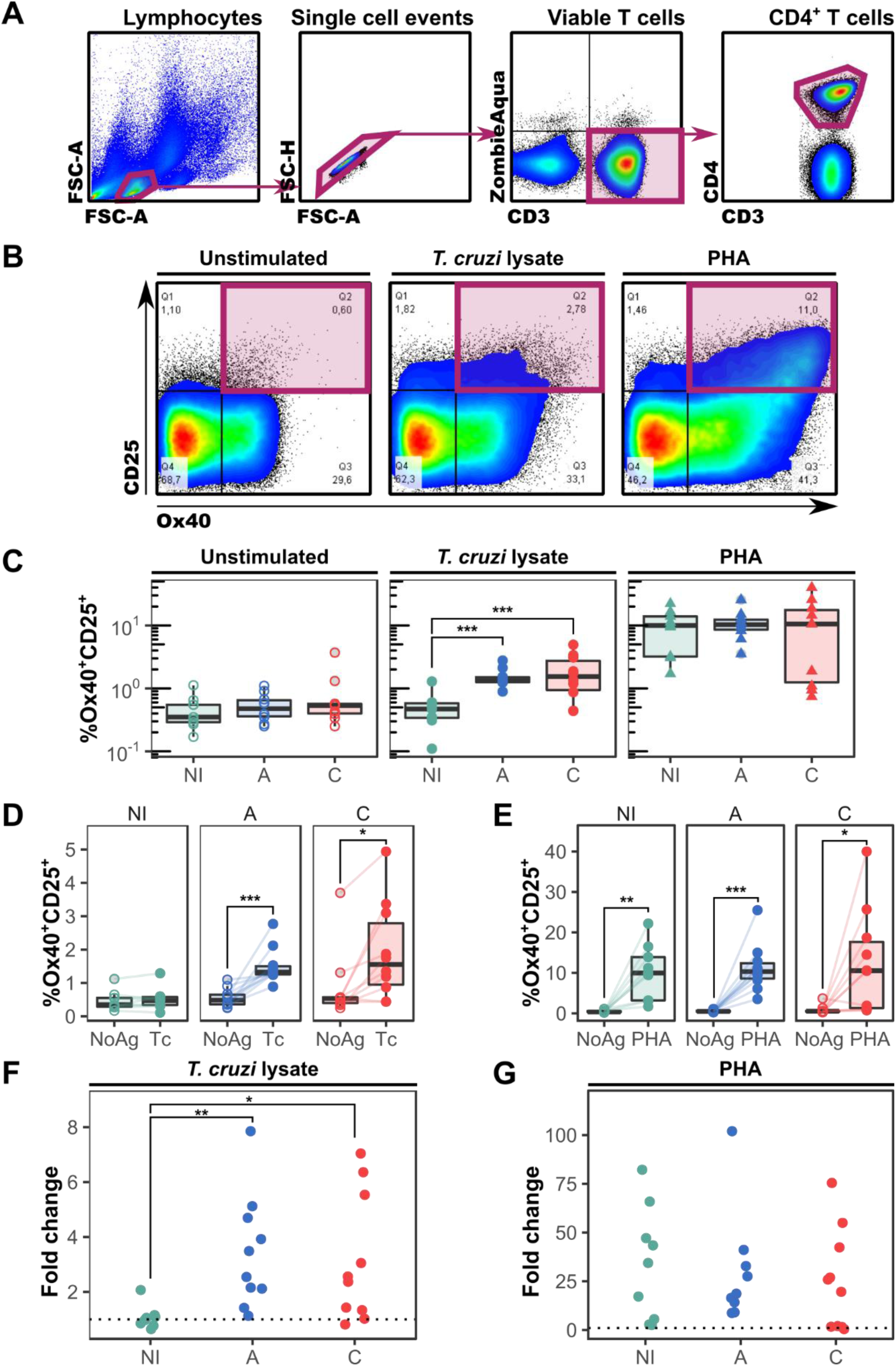
AIM assay with markers Ox40 and CD25 is useful to assess CD4^+^ T cell activation against *T. cruzi* antigens. **A.** Schematic representation of the gating strategy used to analyze CD4^+^ T cell activation. **B.** Representative cytograms from a chronic Chagas patient from group A, under different stimulation conditions. **C.** Differences between groups in the frequency of Ox40^+^CD25^+^ CD4^+^ T cells under different stimulation conditions. **D, E.** Frequency of Ox40^+^CD25^+^ CD4^+^ T cells upon *T. cruzi* lysate (**D**) or PHA (**E**) stimulation, paired by subject. **F, G.** Fold change in the frequency of Ox40^+^CD25^+^ CD4^+^ T cells upon stimulation with parasite lysate (**F**) or PHA (**G**), with respect to the unstimulated condition. *: *p*<0.05; **: *p<*0.01; ***: *p*<0.005. NoAg: unstimulated condition; Tc: *T. cruzi* lysate stimulation.

As shown in **Figure 1C-E**, while the frequency of activated cells was similar between groups both upon non-antigen specific stimulation with PHA (*p*=0.9) or without stimulation (*p*=0.5), statistically significant differences were observed in the frequency of Ox40^+^CD25^+^ CD4^+^ T cells between each of the chronic Chagas patients groups and the non-infected subjects when cells were stimulated with parasitic antigens (*p*=0.003 for A vs. NI, *p*=0.01 for C vs. NI, **Figure 1C**). All groups responded significantly to PHA stimulation in comparison with the unstimulated condition (**Figure 1E**). No difference was observed between groups A and C in response to parasite antigens. In addition, the fold change in the frequency of CD4^+^Ox40^+^CD25^+^ T cells upon stimulation with *T. cruzi* antigens, but not with PHA, was greater for both groups of infected patients than for the NI control group (**Figure 1F, G**).

In conclusion, the AIM method using the marker combination Ox40/CD25 can be used to detect anti-*T. cruzi* response in CD4^+^ T cells from chronic Chagas disease patients, upon *in vitro* stimulation with parasite lysate.

### Chronic Chagas disease patients have increased frequencies of circulating CD4^+^TIGIT^+^ T cells

Next, we assessed the expression of the inhibitory receptor TIGIT on circulating CD4^+^ T cells from Chagas disease patients and control donors. **Figure 2A** represents the expression profile of this receptor in representative donors from each group.

**Figure 2.**
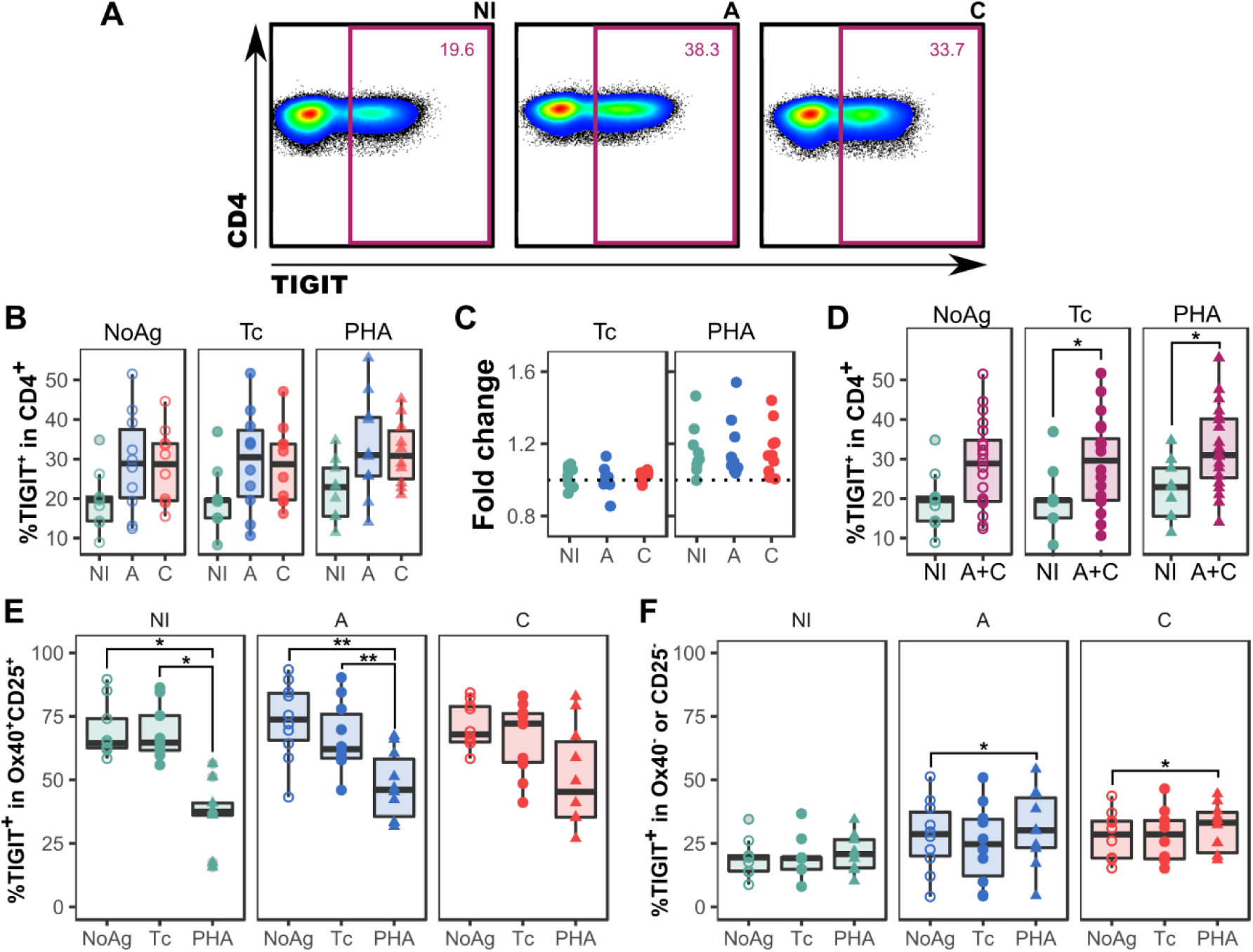
Expression of TIGIT in CD4^+^ T cells from non-infected subjects and chronic Chagas disease patients. **A.** Representative expression patterns in one subject from each group. **B.** Frequency of TIGIT^+^ cells in CD4^+^ T cells for each group, under different stimulation conditions. **C.** Fold change of the frequency of CD4^+^TIGIT^+^ upon parasite lysate or PHA stimulation, relative to that in the unstimulated condition. **D.** Frequency of TIGIT^+^ cells in CD4^+^ T cells, in non-infected subjects and both groups of chronic Chagas disease patients (A+C) collapsed. **E, F.** Frequency of TIGIT^+^ cells in CD4^+^ T cells, for each experimental group, under different stimulation conditions within the activated (Ox40^+^CD25^+^, **E**) and non-activated (Ox40^-^ or CD25^-^, **F**) populations. *: *p*<0.05; **: *p<*0.01; ***: *p*<0.005. NoAg: unstimulated condition; Tc: *T. cruzi* lysate stimulation.

As depicted on **Figure 2B** and **C**, the frequency of TIGIT^+^ cells was not affected in each case by stimulation with *T. cruzi* antigens, nor with PHA. Nonetheless, a striking difference was observed between chronic Chagas disease patients and non-infected individuals. Although statistical significance was not reached when each of the groups were considered separately, the frequency of CD4^+^TIGIT^+^ T cells was significantly higher in chronic Chagas disease patients than in the control group subjects upon stimulation with *T. cruzi* lysate or PHA (*p*=0.047 and *p=*0.021 respectively, **Figure 2D**). In the unstimulated condition, although there is a visible trend with a low *p-*value (*p*=0.059), the difference is, strictly speaking, non-statistically significant.

The expression of TIGIT was also analyzed separately within activated (Ox40^+^CD25^+^) or non-activated (Ox40^-^ or CD25^-^) CD4^+^ T cells. Upon non-specific stimulation with PHA, the frequency of TIGIT-expressing activated CD4^+^ T cells was significantly reduced (*p*>0.01) in comparison with the unstimulated and the lysate conditions, but this was not observed for chronic Chagas patients with cardiac manifestation (**Figure 2E**). When the non-activated cells were analyzed, only subjects from the chronic Chagas patients groups showed an increased frequency of TIGIT^+^ events upon stimulation with PHA as compared with the non-stimulated condition. No other differences were observed regarding the effect of the stimuli on the cells of each of the groups.

### Tim-3-expressing CD4^+^ T cells are more abundant in the circulation of chronic Chagas patients with compromised cardiac function

Since Tim-3 is another inhibitory receptor related to T cell exhaustion, we decided to investigate its possible implications for chronic Chagas disease. The frequency of CD4^+^ T cells expressing Tim-3 was assessed using the same experimental approach described above. The expression profiles for these cells in a representative subject from each group are shown in **Figure 3A**.

**Figure 3.**
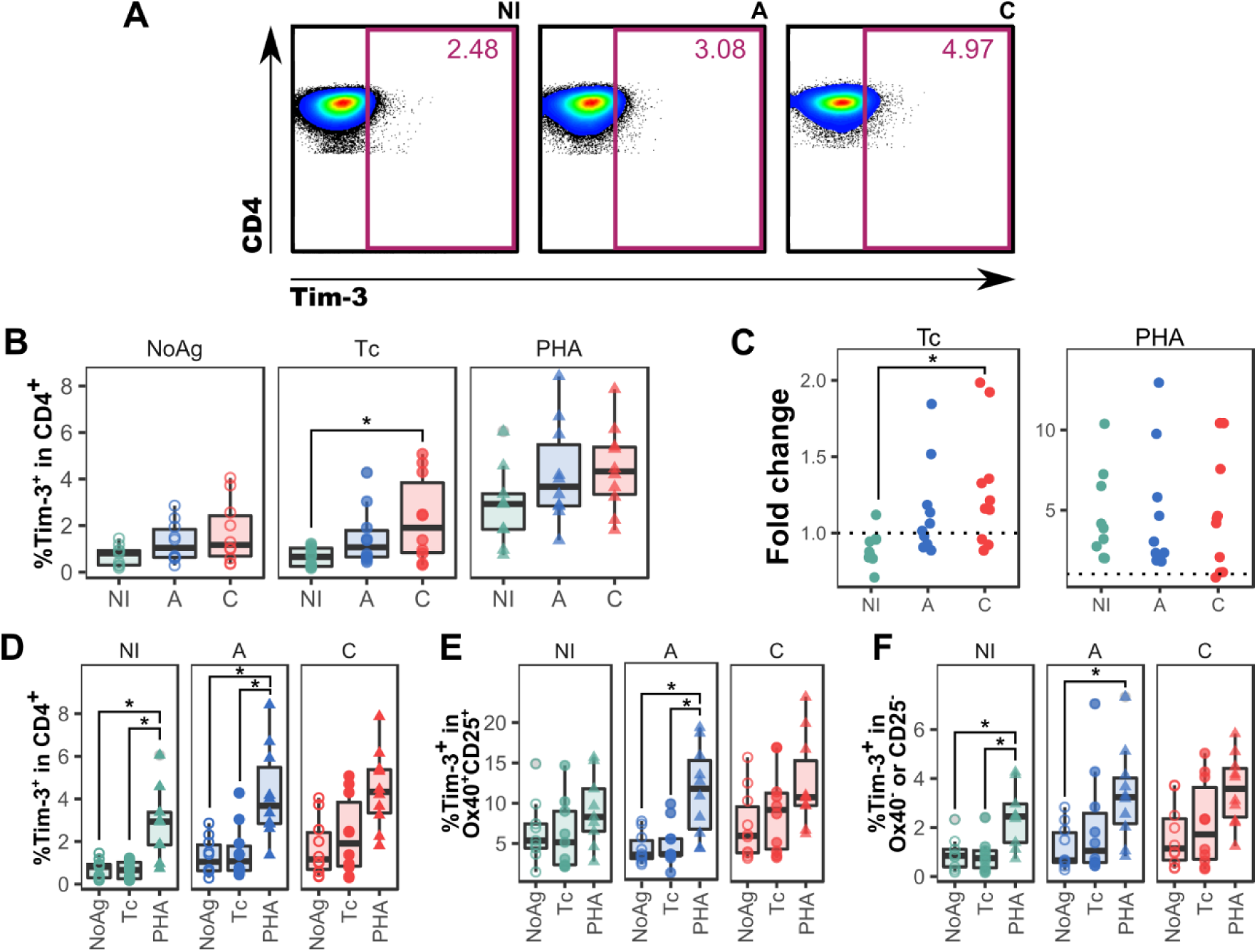
Expression of Tim-3 in CD4^+^ T cells from non-infected subjects and chronic Chagas disease patients. **A.** Representative expression patterns in one subject from each group under the same stimulation condition. **B.** Frequency of Tim-3^+^ cells in CD4^+^ T cells for each group, under different stimulation conditions. **C.** Fold change of the frequency of CD4^+^Tim-3^+^ upon parasite lysate or PHA stimulation, relative to the value in the unstimulated condition. **D-F.** Frequency of Tim-3^+^ cells for each stimulation condition, grouped by experimental groups (NI, A and C), in total (**D**), activated (**E**) and non-activated (**F**) CD4^+^ T cells. *: *p*<0.05; **: *p<*0.01; ***: *p*<0.005. NoAg: unstimulated condition; Tc: *T. cruzi* lysate stimulation.

The results depicted on **Figure 3B** revealed that the frequency of CD4^+^Tim-3^+^ T cells were statistically undistinguishable between groups in absence of antigenic stimulation, or upon non-specific stimulation with PHA. However, exposure to *T. cruzi* lysate unveiled differences in the frequency of such cells, which is higher in subjects from the C group compared to the NI group (*p*=0.04). In fact, the fold increase in the expression of this receptor is greater for chronic Chagas patients in group C than for the other two groups (**Figure 3C**), with the difference being statistically significant only for the comparison between C and NI subjects (*p*∼0.008, medians: 0.91, 1.03 and 1.28 for NI, A and C respectively). This difference was observed separately in both the activated and the non-activated subsets upon encounter with parasite antigens (**Figure 3D**).

Within each group, the abundance of Tim-3^+^ cells was affected by stimulation with PHA, being significantly higher compared to the non-stimulated and the parasite lysate stimulation conditions, in non-infected subjects and patients from group A, but not in patients from group C (**Figure 3D**). Of note, when activated and non-activated CD4^+^ T cells were analyzed separately, the same difference was observed in the non-activated subset for both the A and NI groups, but the A group showed a statistically significant increase in activated Tim-3^+^ CD4^+^ T cells as well (**Figure 3E, F**).

### Lag-3 expression may withdraw CD4+ T cells from activation against T. cruzi antigenic stimulus in chronic Chagas patients without cardiac compromise

The third and last inhibitory receptor we looked into in the context of chronic Chagas disease was Lag-3. The expression profile of this receptor under different stimulation conditions is shown for a representative subject in **Figure 4A**.

**Figure 4.**
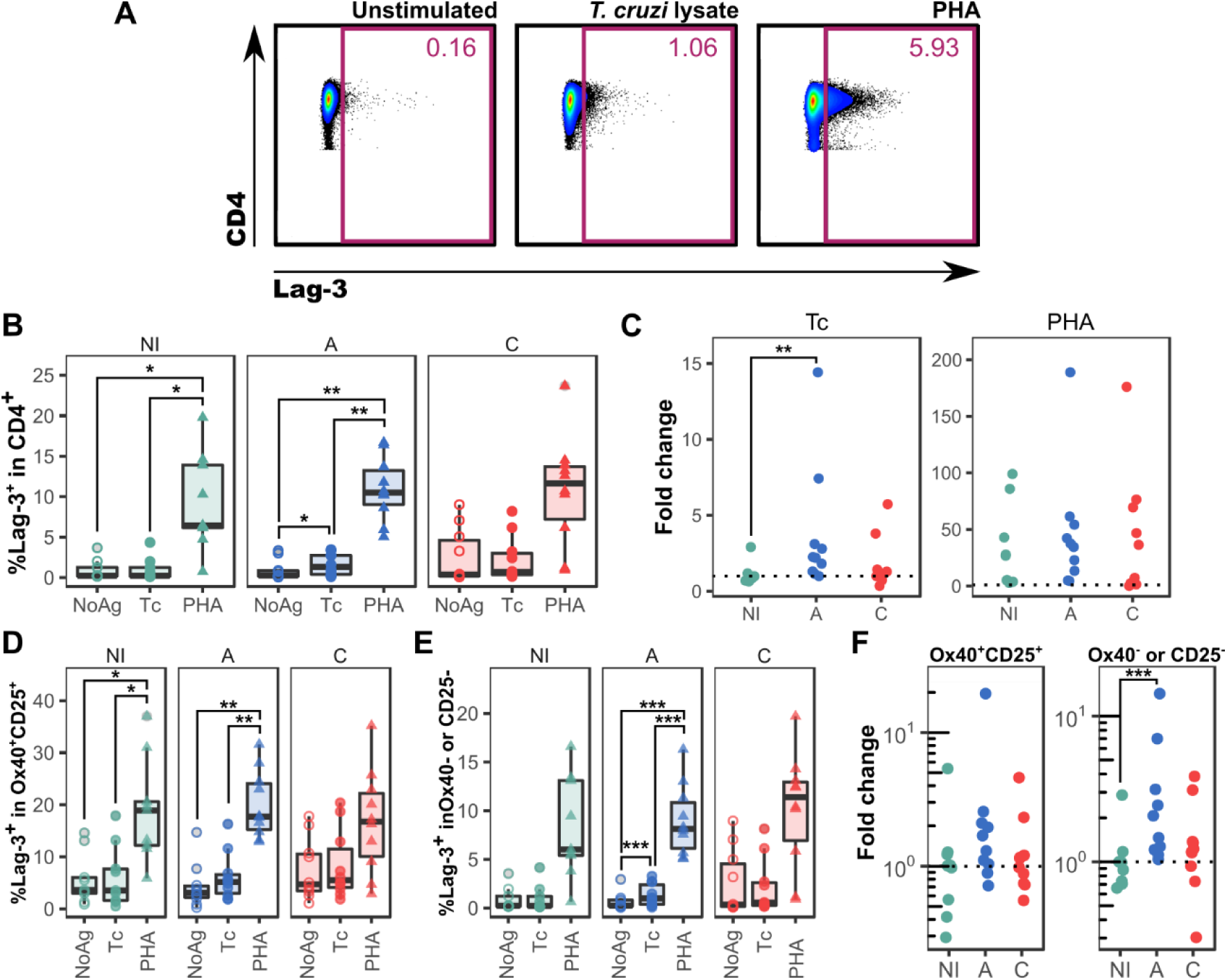
Expression of Lag-3 in CD4^+^ T cells from non-infected subjects and chronic Chagas disease patients. **A.** Representative expression patterns in one subject from group A, under each stimulation condition. **B.** Frequency of Lag-3^+^ cells in CD4^+^ T cells under different stimulation conditions, grouped by experimental groups (NI, A and C). **C.** Fold change of the frequency of CD4^+^Lag-3^+^ upon parasite lysate or PHA stimulation, relative to the value in the unstimulated condition. **D**,**E.** Frequency of Tim-3^+^ cells for each stimulation condition, grouped by experimental groups (NI, A and C), in activated (**D**) and non-activated (**E**) CD4^+^ T cells. **F.** Fold change of the frequency of CD4^+^Lag-3^+^, relative to the value in the unstimulated condition, upon parasite lysate stimulation in activated (Ox40^+^CD25^+^) and non-activated (Ox40^-^ or CD25^-^) CD4^+^ T cells. *: *p*<0.05; **: *p<*0.01; ***: *p*<0.005. NoAg: unstimulated condition; Tc: *T. cruzi* lysate stimulation.

While no statistically significant differences were found in the frequency of CD4^+^Lag-3^+^ T cells between groups in any of the stimulation conditions, this parameter was significantly affected by stimulation with PHA for subjects from groups A (*p*=0.005) and NI (*p*=0.01). The patients in group C showed an increase of these cells, but the difference did not meet the significance cutoff (*p*=0.054). Interestingly, only patients in group A showed an increase in representation of Lag-3^+^ cells upon stimulation with parasite lysate (*p*=0.017), which was also seen as a significantly higher fold change value for this subset upon *T. cruzi* antigens stimulation, compared to that shown by NI subjects (**Figure 4C**, *p*=0.023).

Further subsetting of the CD4^+^ T cells population according to CD25 and Ox40 revealed, remarkably, that the increase in representation of Lag3-expressing cells observed in group A patients was virtually ascribed to non-activated cells (*p*=0.003), while no significant change was observed in the frequency of Lag3^+^ cells within the activated cells gate (*p*=0.068, **Figure 4D, E**). The fold change in size of this population reflects this conclusion (**Figure 4F**).

## Discussion

In the light of the clear link between T cell exhaustion and chronic Chagas disease, the expression profile of inhibitory receptors and their implications for anti-parasite response may help shedding light on the pathogenesis of inflammatory disease associated to *T. cruzi* infection. In this article, we report that TIGIT, Tim-3 and Lag-3 are expressed differently, or they behave in diverging ways upon *in vitro* stimulation, in chronic Chagas disease patients with different degrees of cardiac compromise.

The diversity of response profiles that CD4^+^ T cells may display in reaction to their cognate epitopes poses an obstacle for the determination of overall activation within this subset. Hence, the discovery of a combination of surface markers, addressable by flow cytometry, that may serve to this purpose^12–14^ was a critical development for experimental designs like the one presented herein. Our data showed that in experimental exposure to *T. cruzi* antigens, the combination of markers Ox40 and CD25 clearly signs an activation of CD4^+^ T cells with memory response characteristics, as it is significantly different in chronic Chagas disease patients than in naïve subjects.

TIGIT is an immunomodulatory receptor expressed by T and NK cells, which binds to CD112 and CD155, producing an inhibitory signal that negatively regulates cellular response and IL-12 production from mature dendritic cells while promoting IL-10 secretion^11,20^. In addition, TIGIT^+^ Treg cells have been shown to selectively inhibit Th1 and Th17, but not Th2 cells^21^. In HIV infection, it is co-expressed with PD-1, and may be regarded as an indicator of T cell exhaustion and severity of the infection, as TIGIT^+^ CD4^+^ T cells are more frequent in viral non-controllers than in elite controllers and non-infected individuals^22^. Our results indicate that the frequency of circulating CD4^+^ T cells expressing this surface marker is elevated in chronic Chagas patients from both groups analyzed in this study. It is worth noting that TIGIT expression has been reported to be induced at the event of TCR-signaled activation of CD4^+^ T cells^23,24^. In contrast, in our experimental set up, the frequency of CD4^+^ expressing this marker showed no change upon stimulation with parasite lysate, and a maigre increase with PHA. A tendency is visible in samples from the A group by which the frequency of TIGIT^+^ cells is reduced within the Ox40^+^CD25^+^ subset in the lysate activated condition. This trend, which is evident in the PHA activated condition for all groups, may be associated with a decrease in the representation of Treg cells inside that population in favour of non-regulatory activated T cells. Of note, this was not observed in patients from the C group.

In order to interact with its ligand, galectin 9 (Gal-9), Tim-3 has a mucin domain that is highly glycosylated^11,25^. This is of particular interest in the case of chronic Chagas disease, as *T. cruzi* has been shown to tamper with T cell regulation by altering their surface glycosylation pattern via enzymes of the trans-sialidase superfamily (Marques da Fonseca 2019). Of note, a link between the expression of Gal-9 in myenteric ganglia from chronic Chagas patients and the enteric manifestation of the disease has been proposed^26^. Whether this is also the case in the cardiac forms of chronic Chagas disease, and its possible implications for immune regulation in this pathology via Tim-3 is yet to be investigated. Nonetheless, Lasso et al.^27^ described that this receptor is expressed in higher frequencies of CD8^+^ T cells in chronic Chagas patients than in non-*T. cruzi* infected donors. Our observations indicate that this difference also applies to CD4^+^ T cells. In fact, the frequency of Tim-3^+^ cells within this population increases in chronic Chagas patients but not in non-infected subjects, upon exposure to parasite antigens. In addition, this inhibitory receptor seems to be upregulated in non-activated cells under this condition, despite this difference not being statistically significant.

Lag-3 is a negative regulator of T cell proliferation. It is believed to exert its inhibitory function, not only by competing with the CD4 co-receptor for class II MHC binding in the context of immune synapsis, but also by interacting with proteins from other signaling pathways, like LSECtin^4,11^. Its immune suppressive function has been shown to be subordinated to and synergistic with that of PD-1, and therefore is receiving much attention from the oncotherapy research community^11^. Our results show that, in the context of chronic Chagas disease, the frequency of CD4^+^ T cells expressing Lag-3 is boosted by polyclonal activation with PHA in all of the evaluated groups of donors, but only in group A upon parasite-specific stimulation. An outstanding outcome of our experiments is that this increased frequency of Lag-3^+^ cells occurs within the non-activated subpopulation. This may implicate that this receptor is preventing pathogen-reactive CD4^+^ T cells from triggering an effector response. Furthermore, as this was not observed in cardiac patients, it is tempting to think on a possible participation of Lag-3^+^ T cells-mediated modulation of the immune response in preventing inflammatory pathogenesis. At this point, it should be noted that this inhibitory receptor is expressed on natural and induced regulatory T cells^11^. Also, evidence of an increment on the frequency of regulatory T cells induced by *in vitro* stimulation with *T. cruzi* antigens has been reported previously^28^. Further analysis of our data will let us determine whether the increase in the frequency of Lag-3^+^ cells within the so-termed non-activated population occurs within the CD25^+^ subset, or on the contrary involves CD25^-^ lymphocytes only, which would rule out Tregs as accountable for these changes.

Another scope to be incorporated in future versions of this report is that of combined expression of multiple inhibitory receptors. As opposed of multifunctionality and in the light of the apparent regulatory hierarchy these receptors take part in^4^, the conjoined expression of different inhibitory receptors may imply different stages of immune modulation, which may point at differences between groups of patients and shed light on processes relevant for the, still elusive, pathogenesis of Chagas associated heart disease. On that direction, work is underway to develop a multiparametric analytic pipeline that enables, not only practical analysis of the combined expression of multiple markers, but also the evaluation of changes in the levels of expression (measured as mean fluorescence intensity of staining) across multiple experiments.

Finally, experimental *in vitro* blockade of these receptors will help us to better understand the implications of the observations resulting from de data presented herein at a functional level.

